# A 3D *in vitro* model of the human hepatobiliary junction

**DOI:** 10.1101/2025.07.11.664464

**Authors:** Ashley D. Westerfield, Katarzyna A. Grzelak, Katie Katsuyama, Vardhman Kumar, Bess M. Miller, Joa Yun, Jesse Kirkpatrick, David Mankus, Margaret E. Bisher, Abigail K.R. Lytton-Jean, Z. Gordon Jiang, David D. Lee, Christopher S. Chen, Sangeeta N. Bhatia

## Abstract

Cholestasis, or disruption in bile flow, is a common yet poorly understood feature of many liver diseases and injuries. Despite this, many engineered human tissue models of liver disease fail to recapitulate physiological bile flow. Here, we present a 3D multicellular spheroid-based model of the human hepatobiliary junction, the interface between hepatocytes and cholangiocytes often disrupted in liver disease that is required for directing bile excreted by hepatocytes into the biliary ductal system. Building on advances in organoid and spheroid engineering, we co-aggregate human hepatocytes and intrahepatic cholangiocytes into adult hepatobiliary organoids (aHBOs) that structurally connect and functionally transport bile. aHBOs directionally transport bile from hepatocyte bile canaliculi to cholangiocyte-lined ductules, which we visualize through a high-throughput imaging assay. Hepatobiliary junction formation and bile flow dynamics are quantified over time using fluorescent bile acid analogs and AI-assisted image analysis. When subjected to hypoxia-reoxygenation, aHBOs recapitulate features of biliary dysfunction that mimics the cholestasis and ischemia-reperfusion injury that complicates liver transplant. Our findings suggest that 1) a reversible reduction in hepatocyte canalicular function under hypoxia, followed by 2) selective cholangiocyte death upon reoxygenation, are processes that potentially contribute to biliary dysfunction upon ischemic injury. This human-derived, scalable platform provides a phenotypically-relevant *in vitro* model for dissecting biliary pathophysiology and lays the groundwork for a therapeutic discovery platform for post-transplant ischemic cholangiopathy and other cholestatic liver diseases.

## Introduction

The liver has a highly organized tissue ultrastructure that allows it to perform over 500 different vital functions, of which a crucial function is bile transport [1]. Cholestatic diseases, a class of disorders in which this function is disrupted, account for 14.2% of patients who receive a liver transplant, which is often their only curative treatment option [2]. Because of the shortage of available donor livers, recent progress has been made toward engineering liver tissues that can serve as a replacement or bridge to liver transplant. However, a therapeutic liver graft capable of bile transport requires structural and functional interaction between the two main parenchymal cell types of the liver: hepatocytes and bile duct epithelial cells, also called cholangiocytes.

Both cell types establish and maintain cell polarity, in which the apical side is exposed to bile components that are excreted into the digestive tract, while the basolateral side is exposed to nutrients and oxygen in sinusoidal circulation. Hepatocytes polarize into a multipolar cell, in which their apical surfaces wrap around the cells like a belt [3]. These surfaces, called the bile canaliculi, form a “chicken-wire” network passing between cells within the liver parenchyma, into which active transport of bile acids takes place [4]. Cholangiocytes polarize into monopolar cells that are more typical of simple cuboidal epithelial cells, with one side housing the apical membrane and the other side the basal membrane. These cells line hierarchically larger branching ductal structures of the intrahepatic and extrahepatic bile ducts that form the biliary tree. Bile flows through this biliary tree and exits the liver via the common bile duct, is stored in the gallbladder, and is later released into the small intestine to aid in the digestion of fats and lipids. Despite their differences in polarity, the apical surfaces of hepatocytes and cholangiocytes form a contiguous connection or hepatobiliary junction with each other in order to drain bile out of the liver, referred to as the Canal of Hering. The Canals of Hering functionally connect hepatocyte bile canaliculi to cholangiocyte-lined bile ducts, forming the tiniest ductules structures of the biliary tree. The maintenance of this architecture is crucial to healthy bile synthesis and transport, as a disruption in polarity of either cell type within this junction can lead to cholestatic diseases [5], [6]. Despite the physiological importance of hepatobiliary flow in the Canals of Hering, these features have historically been difficult to characterize in native tissue [7], [8] and as a result, difficult to recapitulate in a 3D engineered tissue system.

Recent advancements in organoid technology have enabled the culture and expansion of human hepatocyte organoids, derived from induced pluripotent stem cells [9], [10], fetal human liver [11], and cryopreserved primary human hepatocytes [12]. Within many of these systems, hepatocytes polarize to transport bile into canaliculi but the bile is unable to drain into any bile duct structures like in the native liver due to a lack of integration with cholangiocytes. At the same time, numerous developments have been made toward culturing human cholangiocytes in 3D organoid systems [13]–[17]. While these cholangiocyte organoids express a biliary gene expression profile and functional phenotype, the ability of these cells to transport and modify bile cannot be properly studied without the synthesis and secretion of bile acids from hepatocytes to cholangiocyte-lined lumina. While coculture systems using rat and mouse cell lines have been developed to replicate this junction [18]–[20], no study has yet assembled the building blocks of mature human hepatocytes and cholangiocytes into a functional human hepatobiliary junction.

In this paper, we leverage both organoid and spheroid technologies to generate adult hepatobiliary organoids (aHBOs), which recreate the minimal functional unit of the human hepatobiliary junction. We scale this process up using a high throughput aggregation-based culture platform, which enables parallel self-assembly of a large number of aHBOs. Using a fluorescently labeled bile acid and an imaging assay, we visualize and quantify bile transport from bile canaliculi to cholangiocyte-lined ductules over time and in various culture conditions. Furthermore, we demonstrate the utility of this platform at modeling biliary pathology; specifically, we model ischemic cholangiopathy, a disease characterized by ischemic damage to the bile ducts, often following liver transplant. When conditioned with a hypoxia-reoxygenation scheme that resembles the ischemia-reperfusion timeline of liver transplant, aHBOs recapitulate a dysfunction in bile flow that follows ischemic injury of the bile duct. Being the first system to recapitulate physiological and pathological bile flow within adult human cells in a 3D context, this hepatobiliary organoid model is ideally positioned to enable the discovery of potential interventions for post-transplant biliary dysfunction.

## Results

### Engineered biaggregate liver spheroids promote structural hepatocyte polarity and the formation of functional bile canaliculi

To model the very finest branches of the biliary tree which are comprised of the bile canaliculi formed by the apical surfaces of hepatocytes, we first sought to engineer human liver spheroids with hepatocyte polarity and functional bile transport. Building upon our lab’s previously established method of liver spheroid generation[21], [22], we set out to find the media conditions that enable human hepatocytes to polarize and form bile canaliculi within biaggregate spheroids made of primary human hepatocytes and supporting fibroblasts (Fig. 1a). We then characterized the expression of genes important to hepatocyte polarity and bile acid synthesis. As measured by RT-qPCR, biaggregate liver spheroids express genes important to hepatocyte polarity at the apical (*BSEP, MRP2*), basolateral (*NTCP*), and tight junction (*CDH1*) portions of the membrane, as well as *CYP7A1*, an important gene in the first step of bile acid synthesis (Fig. 1b). We show that our biaggregate spheroids polarize to form bile canaliculi through immunostaining for F-actin (Fig. 1c), which has been shown to localize at the apical surface of hepatocytes [23]. Additionally, we show that bile acid transporters are not only expressed in biaggregates at the mRNA transcript level, but also at the protein level and are localized at the apical membrane with F-actin, highlighting the formation of structural bile canaliculi within the spheroids (Fig. 1d,e). We further confirm the presence of bile canaliculi within our biaggregate spheroids using scanning transmission electron microscopy (STEM), which shows the formation of lumen structures between two (Fig. 1F, left) or three (Fig. 1F, right) different hepatocytes.

**Figure 1.**
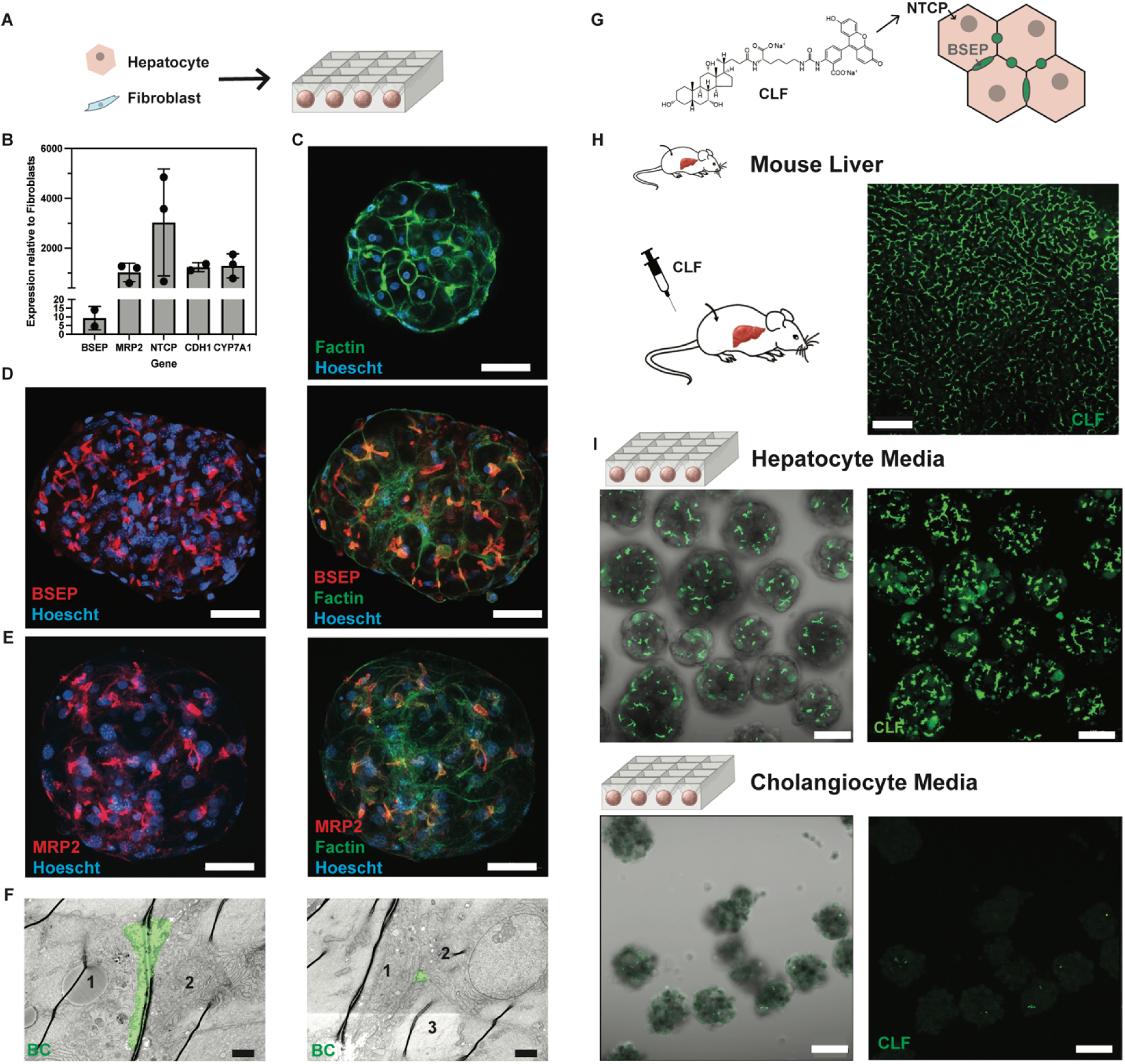
Engineered human biaggregate spheroids polarize to form bile canaliculi that secrete and transport bile acids. A. Schematic depicting the formation of biaggregate spheroids, by incubating a suspension of primary human hepatocytes and supporting fibroblasts in custom V-bottom aggrewells. B. RNA expression of genes involved in hepatocyte polarity and bile acid synthesis, as measured by RT-qPCR in biaggregate spheroids after 3 days in culture. Protein expression of F-actin (C), BSEP (D), and MRP2 (E) in biaggregate spheroids in 3D confocal microscope images of hepatocyte spheroids after 3 days in culture, scale bars = 50um. F. Representative STEM images of bile canaliculi (pseudocolored in green) in biaggregate spheroids after 3 days in culture, scale bars = 2um. G. Schematic depicting the uptake and secretion of cholyl-lysyl fluorescein (CLF) by polarized hepatocytes. H. CLF secretion in bile canaliculi of mouse liver tissue after intravenous delivery, scale bar = 100um. I. CLF secretion in bile canaliculi of biaggregate spheroids after 3 days of culture in hepatocyte media (top) and cholangiocyte media (bottom), scale bars = 100um.

We next established a functional readout of bile acid transport within our engineered biaggregate spheroids. To do this, we adapted an assay using cholyl-lysyl fluorescein (CLF), a fluorescently-labeled bile acid with an image-based readout to visualize bile acid transport to the canaliculi in our engineered biaggregate spheroids (Fig. 1g). This assay has previously been used to visualize bile transport to bile canaliculi in mouse liver tissue [24], in polarized rat hepatocytes within *in vitro* culture platforms [25], and in Canals of Hering-like connections between murine hepatocytes and cholangiocytes [19]. We showed that CLF localizes to the bile canaliculi network in a mouse liver upon intravenous administration of CLF in a saline solution (Fig. 1h). Upon performing the assay in the engineered biaggregate spheroids, we found the CLF to be transported into luminal structures that resemble the size and morphology of bile canaliculi observed in native liver tissue (Fig. 1i). Next, we engineered media conditions to promote the function of these bile canaliculi in biaggregate liver spheroids. We found that media components used in standard hepatocyte medium, such as insulin-transferrin-selenium, fetal bovine serum, and dexamethasone, were important for canaliculi formation. In addition, we found that the inclusion of epidermal growth factor (EGF) increased bile canaliculi number, length, and width (Supp. Fig. 1).

### Hepatocyte polarity and bile canalicular function increases in engineered biaggregate spheroids over time

In order to further characterize bile canalicular formation and function in our engineered biaggregate spheroids, we cultured the biaggregate spheroids over a period of eight days and performed the CLF assay longitudinally (Fig. 2a, b). We developed an image analysis pipeline to measure the number, length, and width of bile canaliculi based on CLF signal in biaggregate spheroids (Fig. 2b, c). Quantification of several spheroids revealed that bile canaliculi increased in number, length, and width over time in culture (Fig. 2c). We also confirmed that our spheroids produced albumin, an important indicator of hepatocyte maturity and function, in these media conditions (Fig. 2d). We also observed the steady increase in total bile acid (TBA) concentration in the media (Fig. 2e), indicating that biaggregate spheroids produce and secrete bile acids into the culture media. To further characterize the composition of bile acids in our samples, we performed ultra-performance liquid chromatography-tandem mass spectrometry (UPLC-MS/MS) on media from our spheroids to detect a panel of 12 different primary bile acids typically found in human bile (Supp. Fig. 2). We found that the majority of the bile acids detected were conjugated with taurine or glycine, which typically indicates a healthy bile acid composition, as conjugated bile acids are typically more water soluble than unconjugated bile acids [26].

**Figure 2.**
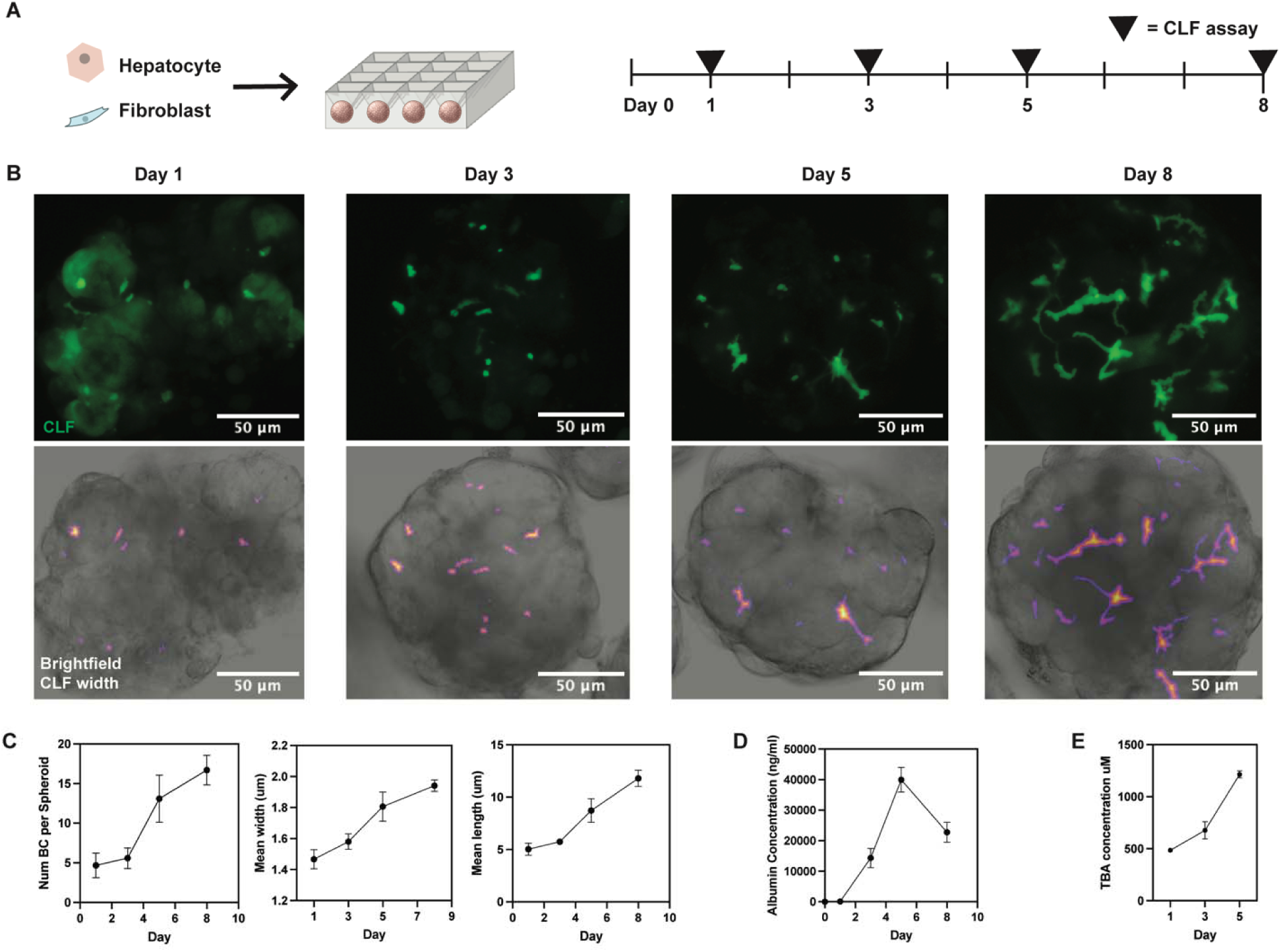
Characterization of bile acid synthesis and transport in engineered biaggregate spheroids. A. Schematic depicting timeline of biaggregate spheroid culture and CLF assay timepoints. B. Representative CLF (top) and width-coded (bottom) images of biaggregate spheroids over time, scale bars = 50um. C. Quantification of number (left), mean width (middle), and mean length (right) of bile canaliculi in biaggregate spheroids over time. D. Albumin secreted into media by biaggregate spheroids over time. E. Total bile acids secreted into media by biaggregate spheroids over time.

### Engineered adult hepatobiliary organoids (aHBOs) recapitulate physiological bile transport function

After constructing a model system that structurally and functionally recapitulates the smallest elements of the biliary system, we set out to add the next element of the hepatobiliary junction: intrahepatic ductules that functionally connect to bile canaliculi. To do this, we isolated intrahepatic cholangiocytes from adult human liver biopsy tissue, establishing a reliable source of human intrahepatic cholangiocytes. We cultured and expanded these cells as organoids in 3D Matrigel culture (Supp. Fig. 3), as previously described [15], [16]. Next, we sought to identify the co-culture conditions necessary to not only enable the mutual survival of both hepatocytes and cholangiocytes, but also to facilitate the structural and functional connection between these two cell types, thus creating the minimal functional unit of the hepatobiliary system. To this end, we modified our spheroid aggregation system to generate triaggregate spheroids, consisting of primary human hepatocytes, intrahepatic cholangiocytes, and supporting fibroblasts (Fig. 3a). We tested a variety of cell ratios and media conditions (Supp. Fig. 4), and found that a mixed co-culture media and a 2:2:1 ratio of hepatocytes, fibroblasts, and cholangiocytes enabled the formation of adult hepatobiliary organoids (aHBOs). We found, through RT-qPCR, that aHBOs expressed hepatocyte polarity genes at a similar level to biaggregates (Fig. 3b). Staining with F-actin revealed the presence of both bile canaliculi and budding ductule-like structures within the same aHBO, indicating the survival and polarity of both hepatocytes and cholangiocytes within our engineered culture. Further immunostaining revealed that budding ductule structures were positive for cytokeratin-7 (CK7) and E-cadherin, indicating that they were lined by cholangiocytes (Supp. Fig. 5, Fig. 3e), and the bile canaliculi structures were BSEP-positive (Fig. 3d). STEM imaging also confirmed the presence of bile canaliculi and ductules within the same spheroid (Fig. 3f).

**Figure 3.**
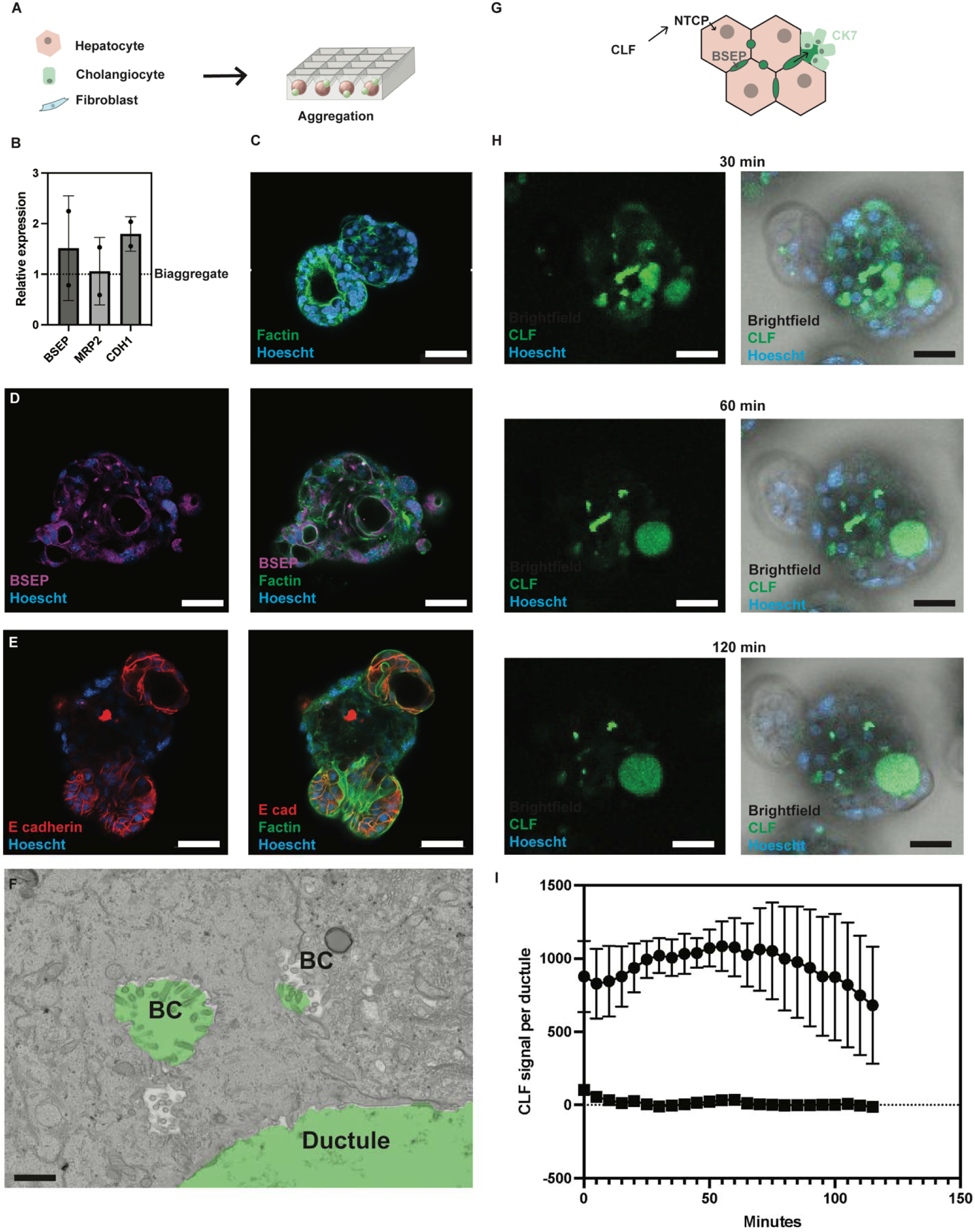
Engineered human adult hepatobiliary organoids (aHBOs) contain hepatobiliary junctions that transport bile from bile canaliculi to intrahepatic bile ductules. A. Schematic depicting the formation of aHBOs with hepatobiliary junctions, by incubating a suspension of primary human hepatocytes, human intrahepatic cholangiocytes, and supporting fibroblasts in custom V-bottom aggrewells. B. RNA expression of genes involved in hepatocyte polarity, as measured by RT-qPCR in aHBOs after 3 days in culture. Protein expression and localization of F-actin (C), BSEP (D), and E cadherin (E) in 3D confocal microscope images of aHBOs after 3 days in culture, scale bars = 50um. F. Representative STEM images of bile canaliculi (BC) and ductule structures (pseudocolored green) in aHBOs after 3 days in culture, scale bar = 1um. G. Schematic depicting the transport of CLF from hepatocyte bile canaliculi to bile ductules. H. Timelapse images of bile acid transport in aHBOs at 30 (top), 60 (middle), and 120 (bottom) minutes after incubation with CLF, scale bars = 50um. I. Quantification of timelapse images of CLF transport in aHBOs in connected ductules (circle) and non-connected ductules (square).

After confirmation that the structural elements of the hepatobiliary junction were present, we sought to characterize the bile transport function within our engineered spheroids. To do this, we used the CLF assay to visualize the connection between bile canaliculi and ductule structures (Fig. 3g). Timelapse imaging of CLF transport in aHBOs demonstrated the initial accumulation of CLF first in bile canaliculi at 30 minutes after incubation (Fig 3h, top). At 60 minutes, the CLF drained from bile canaliculi to connected ductule structures, with the CLF signal within the ductules increasing steadily (Fig. 3h, middle). By 120 minutes, the CLF signal in the bile canaliculi had dramatically decreased with accumulation of signal in connected ductules (Fig. 3h, bottom). This accumulation of CLF signal did not occur in all ductules, implying that the CLF collection was only able to occur in ductules functionally connected to bile canaliculi from which the CLF drained. Non-connected ductules failed to fluoresce with CLF signal throughout the entire 120-minute duration of the experiment (Fig. 3i).

### Characterizing hepatobiliary junction dynamics in aHBOs over time

In our newly developed system, we sought to longitudinally characterize hepatobiliary junction formation and function. To do this, we performed the CLF assay on aHBOs at multiple time points over an 8 day culture period (Fig. 4a). We then implemented an image analysis pipeline that leveraged an AI-guided image segmentation tool to identify CLF-positive ductule structures in a 3D reconstruction of images collected during the CLF assay (Fig. 4b). From this image analysis, we were able to determine the number of CLF-positive ductule structures in each image, normalized by number of aHBOs in the image (Fig. 4c).

**Figure 4.**
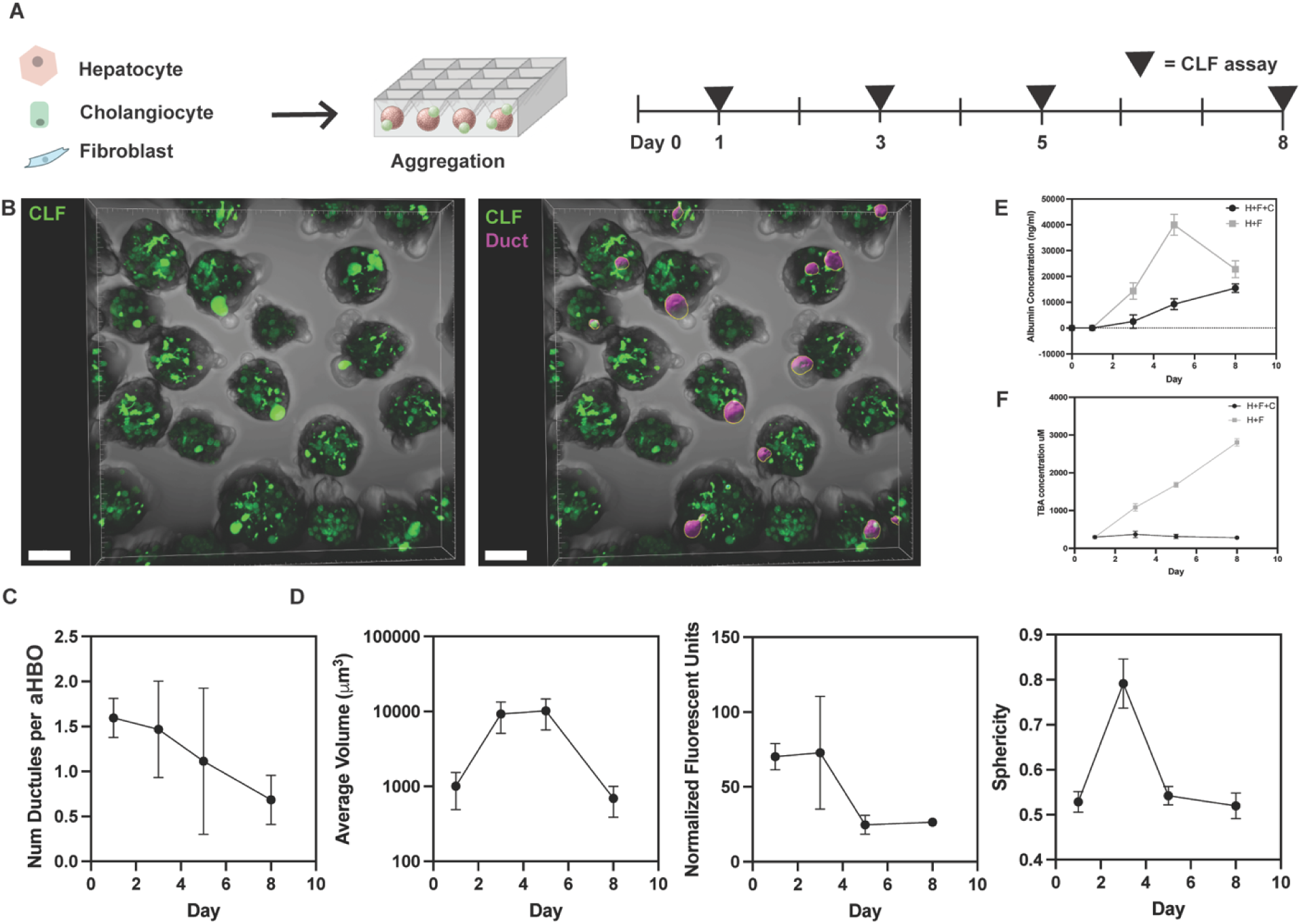
Functional characterization of intrahepatic ductules in engineered aHBOs. A. Schematic depicting timeline of aHBO culture and CLF assay timepoints. B. Representative images of CLF transport from bile canaliculi to bile ductules in aHBOs (left) and segmentation of CLF signal in bile ductules (right, purple), scale bar = 100um. C. Quantification of the number of CLF-positive bile ductules per aHBO over time. D. Quantification of volume (left), fluorescence intensity (middle), and sphericity (right) of CLF-positive bile ductules in aHBOs over time. E. Albumin secreted into media by aHBOs (black) and biaggregate spheroids (gray) over time. F. Total bile acids secreted into media by aHBOs (black) and biaggregate (gray) spheroids over time.

To determine how well these connections maintained their function over time, we measured volume and fluorescence intensity as proxy measurements of how much CLF accumulated in the ductules over time. While the volume of these ductules peaked in size between days 3 and 5 in culture (Fig. 4d, left), the fluorescence intensity decreased but still showed positive CLF signal out to 8 days in culture (Fig. 4d, middle), indicating that the hepatobiliary junctions maintained function over this entire culture period. We also tracked sphericity as a way to quantitatively analyze ductule morphology over time. The ductules were most spherical after 3 days in culture, and while they became slightly less spherical post 5 days in culture, they still maintained function by accumulating CLF (Fig. 4d, right). Taken together, these results show that hepatobiliary junction function peaks in aHBOs around 3 days in culture, but can be sustained up to 8 days in culture.

Lastly, we measured albumin and bile acid synthesis functions in aHBOs, by performing an albumin ELISA and a total bile acid (TBA) assay on media collected from these cultures at various timepoints. We show that aHBOs consistently produce albumin over 8 days in culture (Fig. 4e). However, we see that TBA concentration in media collected from aHBOs is much less than that collected from biaggregates (Fig. 4f). This finding suggests that bile acids may be accumulating in the ductules of the aHBOs, an indication of physiological intrahepatic biliary function. These results demonstrate that aHBOs transport bile in a way that recapitulates intrahepatic bile duct function in the native liver.

### Hypoxia-reoxygenation disrupts bile flow within aHBOs, generating a functional model of biliary dysfunction in post-transplant ischemic cholangiopathy

We next aimed to perturb hepatobiliary function in aHBOs in order to model an ischemic injury to the bile ducts characterized by biliary strictures, known as ischemic cholangiopathy. The mechanism behind biliary strictures is mostly unknown, but is thought to be a result of the ischemia-reperfusion of the bile ducts that commonly complicates liver transplantation [27]. Ischemic cholangiopathy causes delayed liver graft function, and sometimes graft loss, representing a common and debilitating complication in liver transplant that, to this day, has no effective therapy. As such, developing models that capture important aspects of this disease phenotype, including hepatobiliary architecture and bile flow, may potentially contribute to the treatment landscape.

While transport timelines for donor livers can vary widely from patient to patient, the average ischemia time for a donor liver is around 6 hours, during which time the liver is cut off from blood supply before being reconnected to the organ recipient’s systemic circulation [28]. To recapitulate the typical ischemia-reperfusion process that occurs during liver transplantation, after 3 days in normoxic (21% oxygen tension) culture, we subjected aHBOs to 6 hours of hypoxia (2% oxygen tension) and then reoxygenated the spheroids for 24 hours. We conducted the CLF assay in normoxia, after 6 hours of hypoxia, and after 24 hours of reoxygenation (Fig. 5a). We then detected the CLF-positive ductules at these timepoints (Fig. 5b). Under hypoxic conditions, fewer CLF-positive ductule structures were detected. This number was somewhat rescued during reoxygenation, although not quite to the level seen under normoxia (Fig. 5c, top).

**Figure 5.**
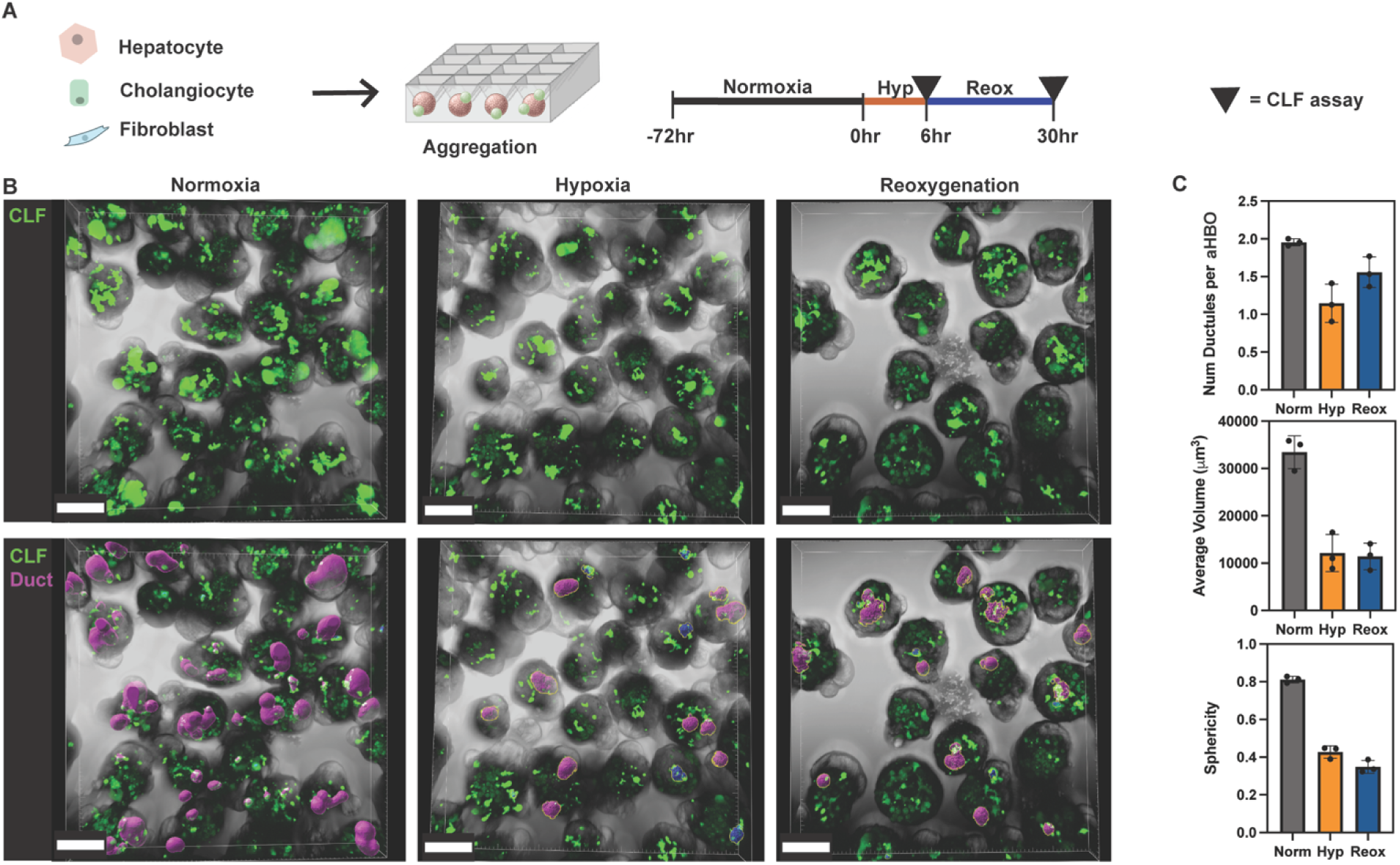
Hypoxia-reoxygenation disrupts bile flow within engineered aHBOs. A. Schematic depicting timeline of hypoxia-reoxygenation conditioning of aHBOs and CLF assay timepoints. B. Representative images of CLF transport from bile canaliculi to bile ductules in aHBOs in normoxic (top left), hypoxic (top middle), and reoxygenated (top right) conditions, and segmentation of CLF signal in bile ductules (bottom, purple), scale bar = 100um. C. Quantification of number (top), volume (middle), and sphericity (bottom) of CLF-positive ductule structures in each condition.

Notably, under hypoxic conditions, CLF-positive ductules have a significantly decreased volume (Fig. 5c, middle) and sphericity (Fig. 5c, bottom), and these features are not recovered to pre-hypoxic levels upon reoxygenation. Taken together, the presence of smaller, less spherical, and more irregularly shaped ductules under hypoxia-reoxygenation suggests that this conditioning causes a volumetric collapse of the ductules, which morphologically resembles the biliary stricturing phenotype seen in ischemic cholangiopathy [29].

### Reversible bile canalicular transport decline under hypoxia is followed by loss of cholangiocyte viability under reoxygenation of aHBOs

Upon observation of biliary dysfunction in aHBOs under hypoxia-reoxygenation conditioning, we sought to conduct this conditioning on the individual cell types in aHBOs in order to understand the processes potentially contributing to this phenotype. First, we aimed to elucidate the effects of hypoxia on CLF transport by hepatocytes into bile canaliculi. To do this, we conditioned our biaggregate spheroids in hypoxia over the course of 3 days and conducted the CLF assay each day (Fig. 6a). CLF signal in bile canaliculi decreased in number, width, and length over the course of three days (Fig. 6b, c), demonstrating that hepatocyte polarity and bile transport function decreases under hypoxic conditions. To determine whether hepatocyte polarity was recoverable upon reoxygenation, we exposed biaggregate spheroids to the same hypoxia-reoxygenation conditioning regimen that induced ischemic cholangiopathy phenotype in aHBOs and conducted the CLF assay in the biaggregate spheroids (Fig. 6d). Under hypoxia, the number of bile canaliculi stayed the same, but the width and length decreased, indicating subtle yet detectable decreases in bile canalicular function (Fig 6e, f). This decrease was completely recovered upon reoxygenation (Fig. 6f), indicating that canalicular transport of bile can be recovered after 6 hours of hypoxia in aHBOs.

**Figure 6.**
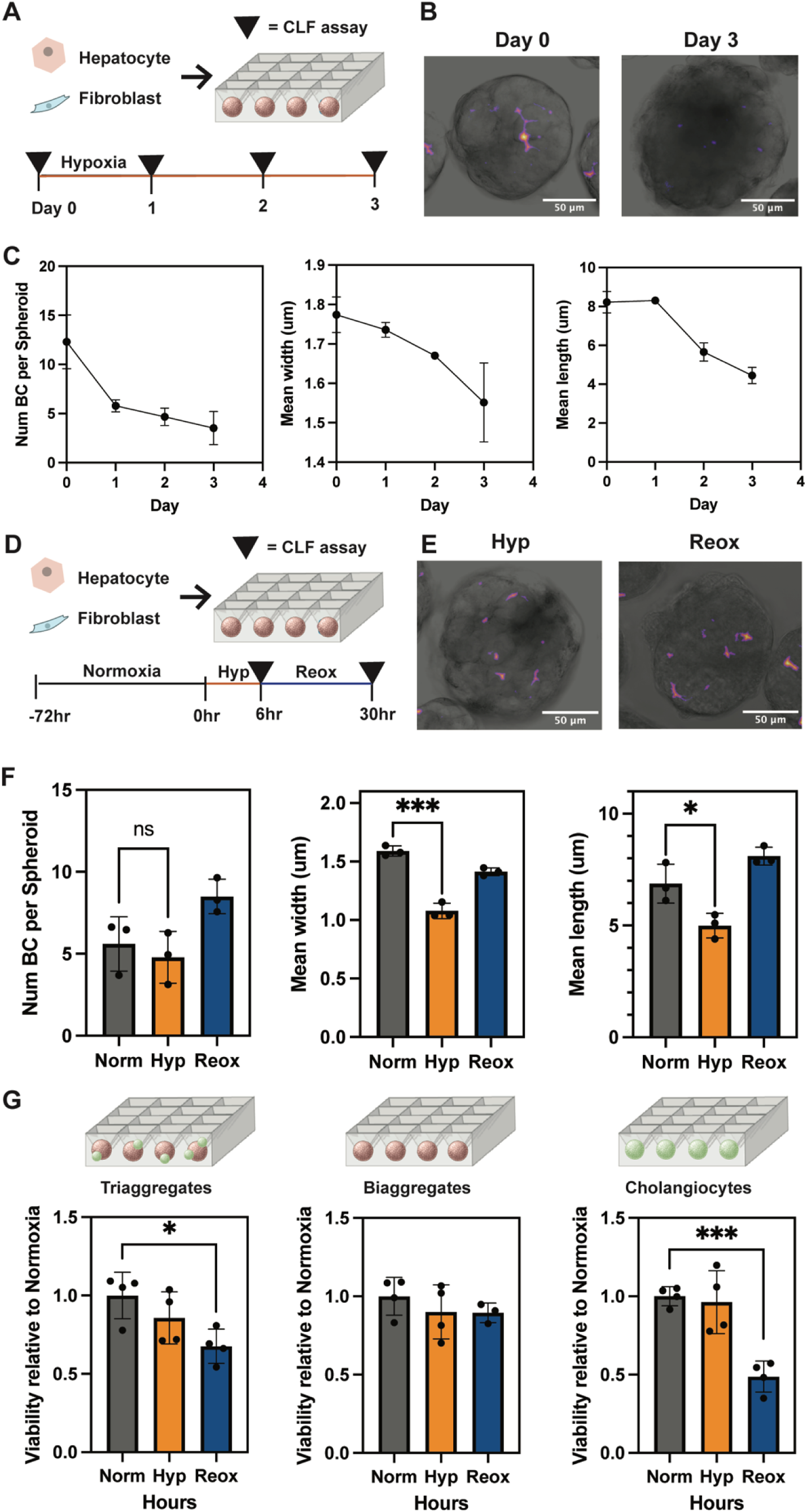
Hypoxia disrupts bile canalicular transport while reoxygenation triggers cholangiocyte-specific cell death in engineered spheroids. A. Biaggregate spheroids were cultured in hypoxia for 3 days, with CLF assay performed on each day. B. Representative images of bile canaliculi in biaggregate spheroids in normoxia (left) and after 3 days of hypoxia (right), scale bars = 50um. C. Quantification of bile canaliculi number (left), width (middle), and length (right) over time in hypoxia. D. Biaggregate spheroids were cultured in hypoxia for 6 hours and then reoxygenated for 24 hours. E. Representative images of bile canaliculi in biaggregate spheroids after 6 hours of hypoxia (left) and 24 hours reoxygenation (right), scale bars = 50um. F. Quantification of bile canaliculi number (left), width (middle), and length (right) in hypoxia-reoxygenation conditioning. G. Viability of aHBOs (left), biaggregate spheroids (middle), and cholangiocyte spheroids (right) during hypoxia-reoxygenation conditioning.

We then wanted to test the effect of hypoxia-reoxygenation on cell viability within aHBOs. We observed a slight decline in cell viability in aHBOs under hypoxia, and this decline was further exaggerated under reoxygenation (Fig. 6g, left). In our biaggregate spheroids, cell viability was unchanged in hypoxia and reoxygenation (Fig. 6g, middle), while spheroids consisting of only cholangiocytes significantly decreased in viability after hypoxia-reoxygenation (Fig. 6g, right). This suggests that the decrease in cell viability can be attributed to the cholangiocytes present in aHBOs. Taken together, these findings demonstrate two processes that contribute to biliary dysfunction after ischemia-reperfusion: The 1) reversible decrease in hepatocyte bile canalicular function under hypoxia coupled with 2) increased cholangiocyte-specific death under reoxygenation leads to a volumetric collapse of ductules in our hepatobiliary units (Fig. 7). Through this work, we demonstrate that aHBOs not only recapitulate bile flow within a healthy hepatobiliary unit, but can also be used to elucidate the mechanisms of clinically-relevant hepatobiliary diseases, as seen by our case study in ischemic cholangiopathy.

**Figure 7.**
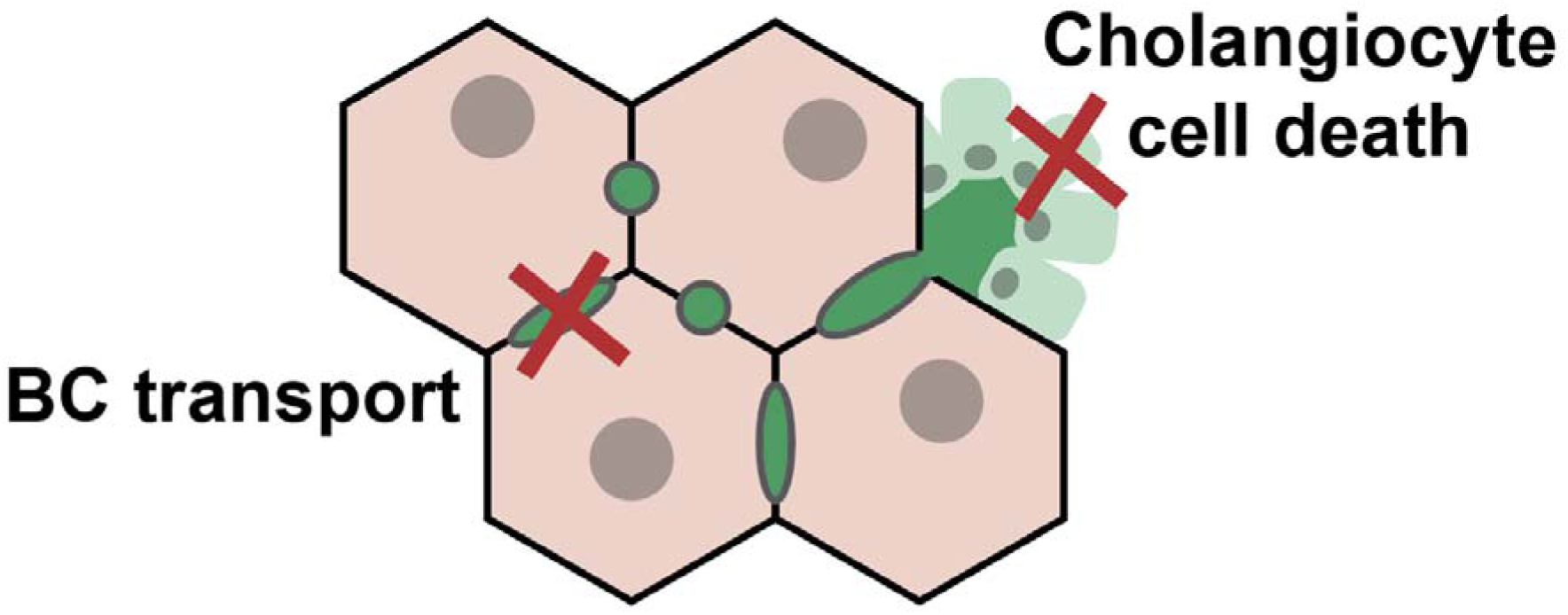
Two potential processes contributing to biliary dysfunction in post-transplant ischemic cholangiopathy.

## Discussion

In this paper, we engineered a 3D *in vitro* model of the human hepatobiliary junction in health and disease, by leveraging a combination of spheroid aggregation and intrahepatic cholangiocyte organoid technology. Specifically, we optimized the culture conditions in which primary human hepatocytes establish polarity and transport bile acids into bile canaliculi within a spheroid context. To visualize and quantify this feature, we developed an image-based assay for high-throughput quantification of bile transport into canaliculi. After establishing a primary human intrahepatic cholangiocyte organoid line, we incorporated these cells into our spheroid platform to engineer aHBOs, which contain a functional hepatobiliary junction between hepatocytes and cholangiocytes. We next adapted our image-based assay to characterize and quantify junction dynamics in aHBOs over time. We then observed bile transport in aHBOs under conditions of hypoxia and reoxygenation, which faithfully recapitulates a phenotypic change seen in post-transplant ischemic cholangiopathy. Recent studies have demonstrated that intrahepatic cholangiocytes undergo intrinsic apoptosis when stressed by hypoxia, followed by extrinsic apoptosis and necroptosis upon reoxygenation, often coupled with upregulation of epithelial to mesenchymal transition (EMT)-associated markers and disintegration of the actin skeleton [30], [31]. While these findings elucidate the cholangiocyte-specific mechanisms of ischemic injury in a monoculture context, the causes and consequences of biliary dysfunction in ischemic cholangiopathy are intricately linked to bile flow [32], [33], which requires a model system that incorporates a functional hepatobiliary junction to study. In this work, we develop such a model system, and build upon recent findings by studying the effect of hypoxia-reoxygenation conditioning in the context of bile transport in a 3D multicellular environment. aHBOs serve as a platform for generating hypotheses regarding the potential processes that contribute to biliary dysfunction in the setting of liver transplantation. These hypotheses include changes in intraluminal pressure caused by the reversible reduction in canalicular flow in hypoxia, as well as compromises in barrier function triggered by cholangiocyte apoptosis in reoxygenation. These hypotheses provide a basis for several future studies.

One limitation of our system is the use of a non-physiological ratio of cholangiocytes to hepatocytes; while our system incorporates a 2:1 ratio of hepatocytes to cholangiocytes, the native liver has much fewer cholangiocytes, with a ratio closer to 16:1 [34]. We justify this design choice to maximize the number of hepatobiliary junctions generated by our platform, such that perturbations to this junction can be observed and averaged over many replicates. In fact, the high-throughput nature of our engineered model system opens the door to a plethora of future screen-based studies. Because the CLF assay serves as a high-throughput, image-based functional readout for connected vs. unconnected ductules, functional genomics screens can be used to elucidate the genetic basis of hierarchical bile transport. Additionally, aHBOs can potentially be used in high-throughput compound screening to develop therapeutics that prevent or reverse biliary dysfunction in patients with ischemic cholangiopathy or, more broadly, patients with a range of cholestatic diseases.

While the use of murine fibroblasts in our spheroid aggregation renders our system chimeric, the cell types that make up the hepatobiliary junction are human-derived. Our lab has previously shown that fibroblasts are essential for aggregation support [35]. We strategically chose a murine fibroblast source so that using human primers in our bulk gene expression analyses (RT-qPCR and bulk RNA sequencing) would reveal gene expression of only the human hepatobiliary cell types within this system, without noise from the fibroblasts. Recent studies have pointed to the role of fibroblasts in biliary fibrosis, by generating an *in vitro* murine liver model in which tuning the ratio of portal fibroblasts induces biliary fibrosis independently of an immune compartment [36]. Future studies could involve incorporating human portal fibroblasts at different ratios to observe changes to hepatobiliary junction in a fibrotic context. Besides fibroblasts, immune cells of the liver microenvironment also contribute to several liver and bile duct diseases. One recent advance in intestinal and liver organoid technology is the integration of macrophages [37], [38]. While these models have focused on viral infection, similar techniques to incorporate an immune component into aHBOs could facilitate future studies in modeling autoimmune cholestatic diseases in which immune cells play a key role, such as primary biliary cholangitis.

Our lab has previously developed liver-on-chip models, which incorporate vascular [39] and bile duct [40] function. Future studies could involve incorporating aHBOs into a microfluidic device-based system that recapitulates the entire hepatic sinusoid in a 3D context, in order to model the synthesis, modification, and recirculation of bile from blood to bile duct, which is a process that is crucial in health and disease [26]. Recent engineered models have been developed to incorporate systemic vasculature [39] and bile ducts [41]. It follows that a complex *in vitro* system that incorporates relevant human liver cells and facilitates access to the lumen of blood vessels and bile ducts could enable us to study these processes in further detail.

In addition to modeling human tissues for *in vitro* studies, the field of regenerative medicine has been advancing toward engineering liver tissues with direct therapeutic utility. We and others have previously developed implantable engineered liver tissues that expand and perform important hepatocyte functions *in vivo* [22]. While these tissues demonstrate hepatocyte polarity *in vivo*, they do not form a contiguous bile canalicular network that functionally drains to a cholangiocyte-lined bile duct, which is a crucial function for the future of engineered liver tissues that can serve as a therapy for patients with cholestatic and bile duct diseases. Incorporating aHBOs into an implantable engineered liver tissue could serve as a promising next step toward developing a fully-functional liver tissue replacement.

Overall, we report the development of adult human hepatobiliary organoids that model both physiological and pathological intrahepatic bile duct function.

## Acknowledgements

The authors would like to acknowledge Dr. Luc van der Laan, Dr. Monique Verstegen, Kimberley Ober-Vliegen, Dr. Gilles van Tienderen, Dr. Meritxell Huch, Dr. Irwin Arias, Dr. Andrea McClatchey, and Dr. Kasturi Chakraborty for thoughtful discussions about experimental design. The authors would also like to acknowledge Dr. Susanna Elledge and Dr. Tahoura Samad for editing the manuscript. A.D.W. acknowledges support from National Science Foundation Graduate Research Fellowship. Z.G.J. receives research support from NIAA (R01AA030770). The work was supported in part by the Koch Institute Support (core) Grant P30-CA14051 from the National Cancer Institute, and we thank the Koch Institute’s Robert A. Swanson (1969) Biotechnology Center for technical support, specifically the Peterson (1957) Nanotechnology Materials Core Facility (RRID:SCR_018674). This work was supported by the NIH (EB033821) and the Wellcome Leap HOPE Program. S.N.B. is a Howard Hughes Medical Institute Investigator.

## Methods

### Human intrahepatic cholangiocyte organoid (ICO) initiation and culture

For initiation of organoid cultures, a fresh donor liver biopsy from a 54-year-old male was collected and stored in cold University of Wisconsin solution. Patient gave informed consent for the use of material for research purposes.

A human ICO line was initiated and cultured using an established protocol [13]. In brief, donor liver tissue was minced using a scalpel, then washed twice via centrifugation in William’s E medium supplemented with 50 ng/ml EGF and 10uM Y27632. The cells were then seeded in basement membrane extract (R&D Systems) in multi-well plates. Organoid initiation medium, comprised of ICO expansion media (Table S1) supplemented with 25ng/ml recombinant human Noggin (Peprotech 120-10C), 30% Wnt3a-conditioned media (made in house), and 10uM Y27632 (SelleckChem, S1049), was then added to the culture. After three days of culture in the initiation medium, ICO expansion media was added to the cells. ICOs were grown and passaged in ICO expansion medium as previously described [13], [15] for 3-6 passages before inclusion in aHBOs. For inclusion in aHBOs, ICO cultures were treated with 5U/ml dispase II (Stemcell Technologies, 07913) until basement membrane extract was dissolved, then TrypLE Express Enzyme (Gibco, 12604021) until a single-cell suspension of cholangiocytes was generated.

### Cell culture

Primary cryopreserved human hepatocytes (Thermo Fisher, Lot #4129) were maintained in Hepatocyte Media (see Table S1). 3T3-J2 murine fibroblasts were a kind gift provided by Howard Green (Harvard Medical School) and were cultured in DMEM with 4.5 g/L glucose (Corning, 10-017-CV), 10% fetal bovine serum, and 1% (v/v) penicillin-streptomycin. Upon confluence, fibroblasts were passaged by treatment with 0.25% Trypsin (Corning, 25-050-CI).

### Spheroid and aHBO formation

AggreWell plates containing 400 μm pyramidal microwells were fabricated in-house and prepared as described previously[21]. To generate biaggregate spheroids, cryopreserved human hepatocytes were thawed and mixed with fibroblasts, in a 1:1 ratio of hepatocytes to fibroblasts. The two-cell mixture was then plated in AggreWell plates in hepatocyte media (Table S1) over the indicated period of time. To generate aHBOs, cryopreserved human hepatocytes were thawed and mixed with fibroblasts and single-cellularized ICOs, in a 2:2:1 ratio of hepatocytes to fibroblasts to ICOs. The three-cell mixture was then plated in AggreWells in a coculture media comprised of 50% Hepatocyte Media and 50% Cholangiocyte Media (Table S1).

### Hypoxia and reoxygenation stimulation in aHBOs

After three days of aggregation in coculture medium (Table S1), aHBOs were transferred to an incubator with hypoxic 1-21% oxygen control (Thermo Fisher, Model 3140), with gas settings at 2% O2, 5% CO2 for 6 hours to mimic transplant-related ischemia-reperfusion injury *in vitro*. aHBOs were then cultured in normoxia (21% O2, 5% CO2) for a reoxygenation time of 24 hours.

### RNA isolation and RT-qPCR

To measure gene expression via RT-qPCR, media was first removed then the sample was lysed and homogenized in TRIzol reagent (Invitrogen, 15596026). RNA was isolated via chloroform extraction and further purified with the RNeasy MinElute Cleanup Kit (QIAGEN, 74204). cDNA was synthesized using RevertAid First Strand cDNA Synthesis Kit (Thermo Fisher, K1622), according to the manufacturer’s instructions. RT-qPCR was performed using PowerUp SYBR Green Master Mix (Thermo Fisher, A25742) in a Bio-Rad CFX96 Real-Time PCR Detection System, according to the manufacturer’s instructions. The primers sequences used to detect mRNA levels are listed in Table S2. Relative mRNA quantification was calculated with the Delta-delta Ct method, using *HMBS* as housekeeping gene.

### Immunostaining

For immunostaining, spheroids were fixed in 4% paraformaldehyde in PBS for 30 minutes – 1 hour. Spheroids were then blocked and permeabilized in 3% bovine serum albumin (BSA) with 0.05% Triton X-100 for 3 hours. Spheroids were then incubated in primary antibody diluted in 3% BSA (according to vendor’s recommendation) at 4C overnight. Spheroids were washed 5 times for 15 minutes each in PBS, then incubated in secondary antibodies (1:500 dilution) and Hoechst 33342 (1:2000) in 3% BSA at 4C overnight. Spheroids were washed 5 times for 15 minutes each in PBS, before being mounted on a coverslip with ProLong Diamond Antifade Mountant (Invitrogen, P36961) and imaged.

### Confocal imaging

Images were obtained on a Zeiss LSM 900 confocal microscope with Airyscan super-resolution detector (Carl Zeiss, Jena, Germany), and processed using ZEN software (Carl Zeiss). Immunostained spheroids were immobilized on a coverslip and imaged using a Plan-Apochromat 20x/0.8 objective (Carl Zeiss, 440640-9903-000).

### In vitro CLF assay

Spheroids were collected from AggreWell plates, gently pelleted at 100g, and incubated in media with 5uM cholyl-lysyl fluorescein at 37C for 30 minutes. Spheroids were briefly washed three times in Advanced DMEM/F12 with 1% penicillin-streptomycin, then plated on a coverslip and imaged in suspension. The same objective, laser intensity, and z-stack interval were used for all samples. For each condition, at least three representative fields of view were captured and analyzed.

### In vivo CLF assay

BALB/c mice (Taconic) were anesthetized using isoflurane. 100 ul of a solution of 1mg/kg CLF in sterile saline was injected intravenously and after approximately 3 minutes, mice were sacrificed and the liver was harvested. The explanted liver was sectioned into 100-200um thick sections with a vibratome (Leica, VT1000S), and imaged on a coverslip using a Plan-Apochromat 20x/0.8 objective (Carl Zeiss, 440640-9903-000).

### Scanning Transmission Electron Microscopy (STEM) imaging

Spheroids and aHBOs were fixed in 2.5% glutaraldehyde and 2% paraformaldehyde in 0.1M sodium cacodylate buffer. Heavy metal staining was performed following a reduced osmium and en bloc uranyl acetate protocol followed by dehydration in a graded ethanol series. Propylene oxide was used as a transitional fluid, and organoids were embedded in Embed 812 resin, and allowed to cure at 60°C for 48 hours. The polymerized resin embedded samples were then sectioned into 60 nm thin slices using a Leica UC7 ultramicrotome and collected onto carbon-coated nitrocellulose film copper grids. Imaging was performed on a Zeiss Crossbeam 540 scanning electron microscope using an annular STEM detector at an accelerating voltage of 30kV and probe current of 200pA.

### Biochemical assays

Conditioned media from spheroid cultures was collected every day and stored at −20 °C. Human albumin was quantified using an enzyme-linked immunosorbent assay (ELISA) using a goat anti-human albumin antibody (Bethyl Laboratories) and 3,3′,5,5′-tetramethylbenzidine (TMB, Thermo Fisher).

Total bile acid content was quantified from conditioned media through enzymatic cycling of bile acids in the presence of NADH and a chromophore using a commercially-available kit (Abcam). Absorbance of the reaction’s colorimetric product was read kinetically at 405nm on a plate reader over time, which could then be converted to a bile acid concentration.

### Image analysis & quantification

To identify bile canaliculi, ImageJ was used to quantify CLF signal. A threshold was placed on each image to identify CLF signal specific to the bile canaliculi (and eliminate intracellular CLF signal). For each image, the quantity (count) of bile canaliculi was recorded, and for each bile canaliculus, the width and length were calculated using the local thickness and Feret’s diameter plugins, respectively. The number of spheroids in each image was quantified manually, and the quantity of bile canaliculi was normalized by the number of spheroids in the image, to account for differences in spheroid density across multiple images.

To quantify ductule structures (lined by cholangiocytes), each image was 3D reconstructed using Imaris software (Oxford Instruments). A surface was rendered for the CLF signal for each image, and the surface was segmented using AI image segmentation tool to identify CLF signal specific to the ductule structures. For each image, the quantity of CLF-positive ductule structures was recorded, and for each CLF-positive ductule structure, the volume and sphericity were recorded. The number of spheroids in each image was quantified manually, and the quantity of CLF-positive ductule structures was normalized by the number of spheroids in the image, to account for differences in spheroid density across multiple images.

### Spheroid viability assay

To measure viability of spheroids after hypoxia-reoxygenation conditioning, the CellTiter-Glo 3D Cell Viability Assay (Promega, Madison, USA) was applied according to the manufacturer’s instructions. Briefly, after conditioning, the spheroids were collected from AggreWell plates and seeded into a 96-well flat white-bottom plate. A volume of CellTiter-Glo reagent equal to the volume of the cell culture medium was added and mixed vigorously with the spheroids. The plate was incubated at room temperature for 25 minutes in the dark. The luminescence were measured by plate reader (Tecan Infinite M Plex, 30190085).

### Statistical analysis

All statistical analyses were performed using Prism software (GraphPad Software Inc.). Data are presented as the mean ± standard deviation (SD) from at least three replicates. A comparison between two groups was conducted using the unpaired Student’s t-test. Comparisons between multiple groups were performed using a one-way ANOVA test followed by Tukey’s post hoc test. For each test, p < 0.05 was considered statistically significant. To indicate statistical significance, one asterisk (*) indicates p < 0.05, two asterisks (**) indicates p < 0.01, and three asterisks (***) indicates p < 0.001.

