## Supplementary Information for "A 3D *in vitro* model of the human hepatobiliary junction"

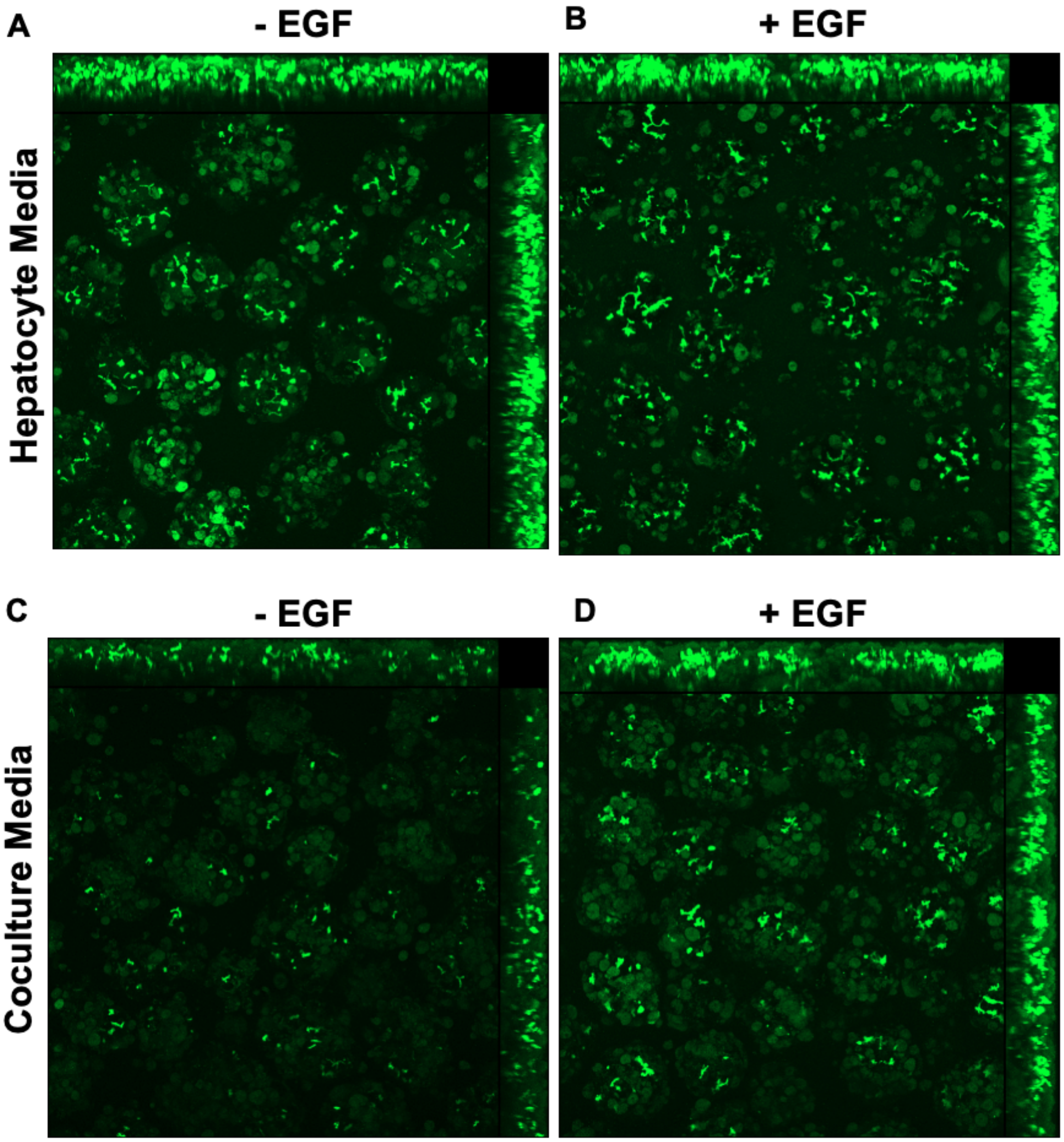


Supplementary Figure 1: EGF improves bile canaliculi formation in biaggregate spheroids in two different media conditions. Representative images of CLF assay on biaggregate spheroids in hepatocyte media without (A) and with (B) 50ng/ml EGF, and in coculture media without (C) and with (D) 50ng/ml EGF.

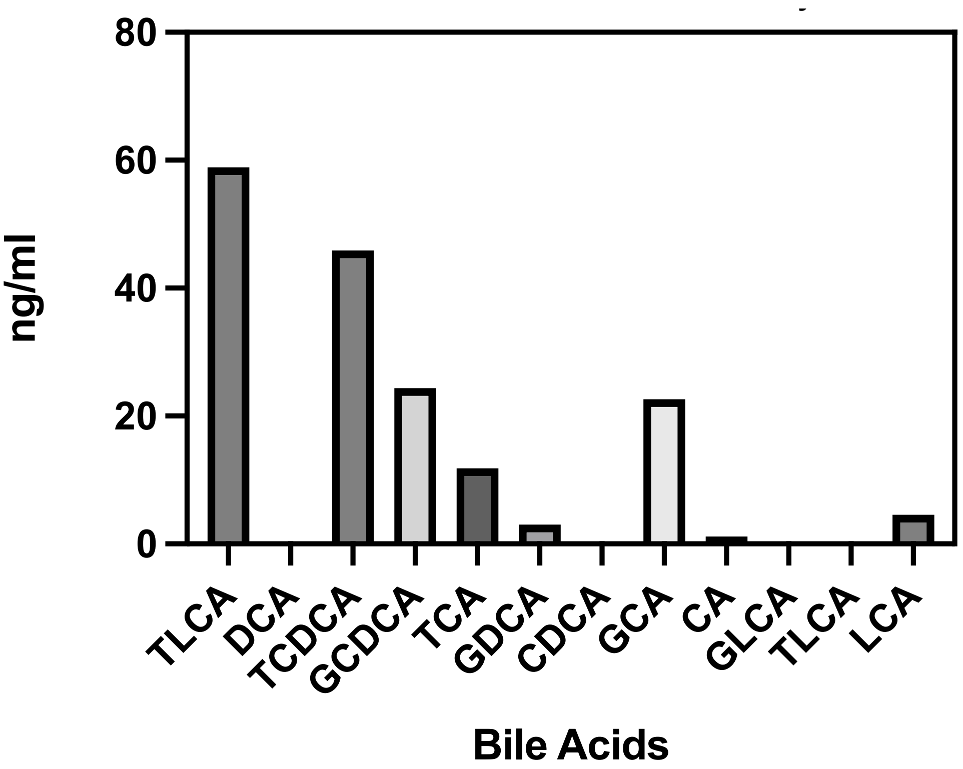


Supplementary Figure 2: LCMS detection of bile acids secreted into media by biaggregate spheroids after 3 days in culture

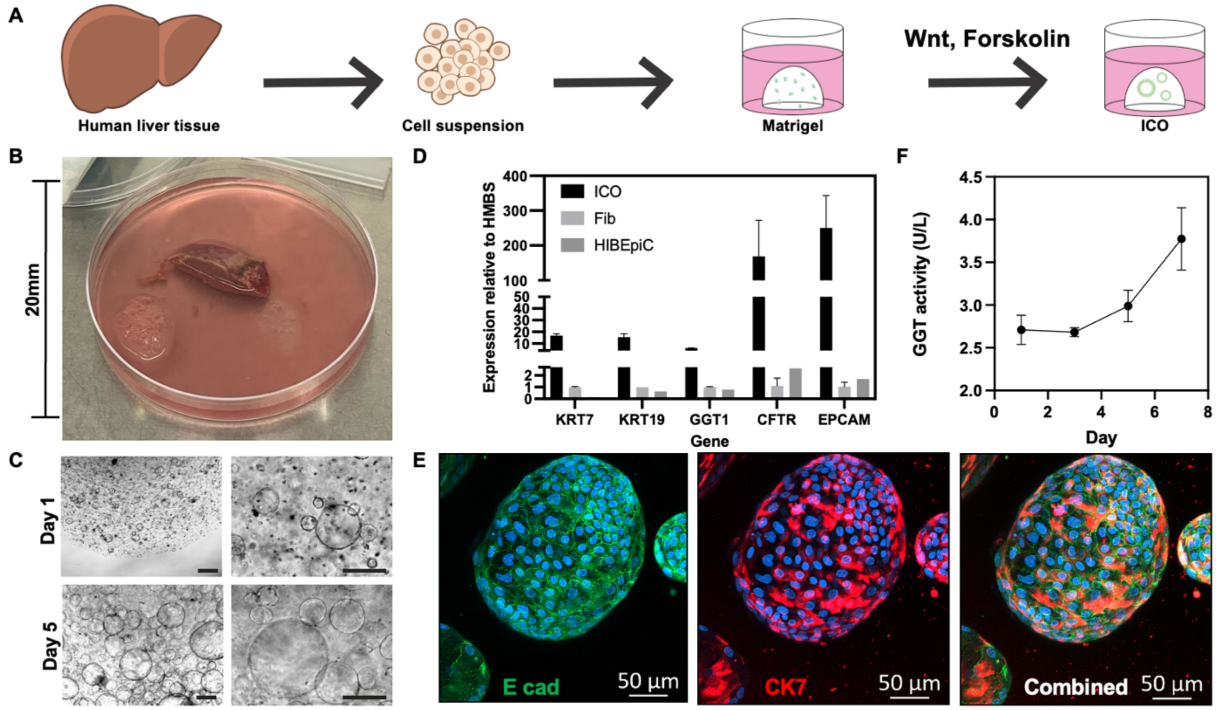


Supplementary Figure 3: Establishing an intrahepatic cholangiocyte organoid (ICO) line from adult human liver biopsy tissue

A. Schematic depicting protocol used to isolate and culture cholangiocytes as organoids from an adult human liver biopsy tissue.
B. Macroscopic image of human liver biopsy tissue (~4mm) in a 20mm Petri dish.
C. Brightfield microscope images of ICO in Matrigel culture, 1 day (top) and 5 days (bottom) after first passage in culture, scale bar = 500um.
D. Gene expression of cholangiocyte specific genes in ICO culture, detected by RT-qPCR. Also graphed are gene expression levels in primary human fibroblasts (Fib) and a commercially-available cholangiocyte cell line (HIBEpiC)
E. Immunofluorescence staining of E cadherin (green) and CK7 (red) in ICOs, scale bar = 50um.
F. GGT activity measured in media collected from ICO culture.

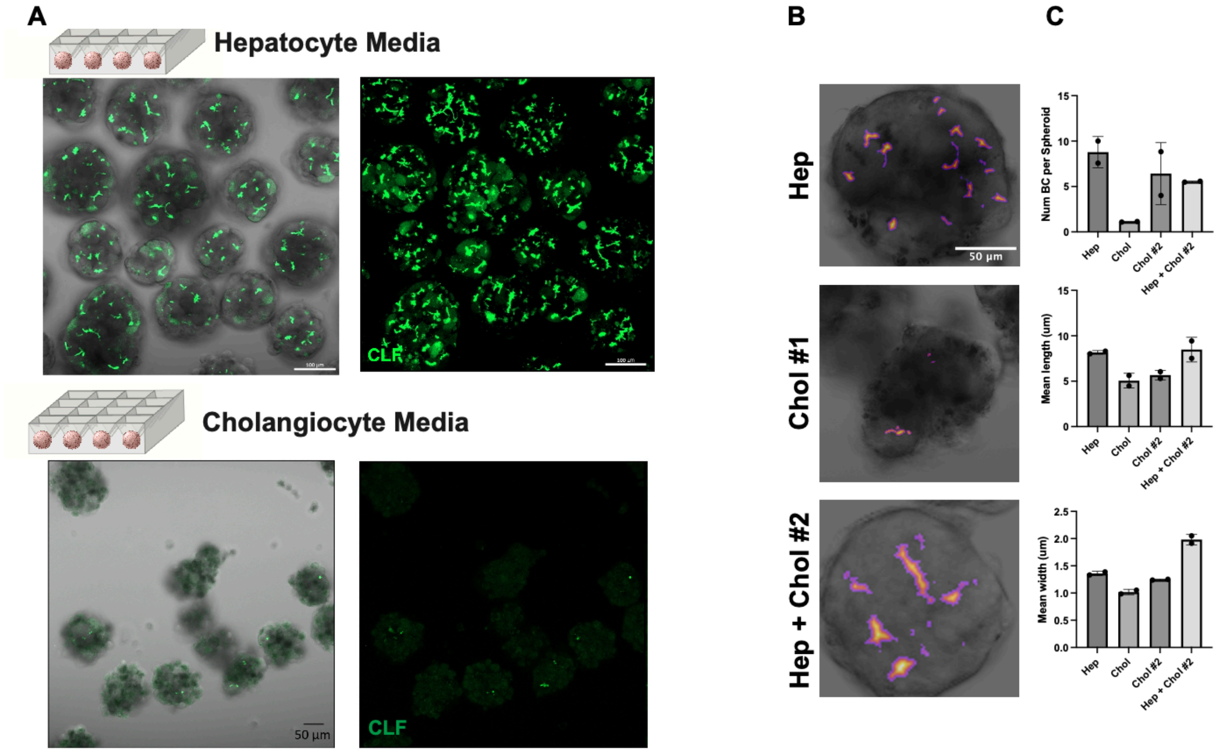


Supplementary Figure 4: Media composition plays a significant role in bile canalicular secretion of CLF in biaggregate spheroids.

(A) CLF secretion in bile canaliculi of 3D hepatocyte spheroids after 3 days of culture in hepatocyte media (top) and cholangiocyte media (bottom).

(B) Representative images width-coded for CLF signal in hepatocyte media (top), cholangiocyte media (middle), and optimized coculture media (bottom).

(C) Quantification of number (top), length (middle), and width (bottom) of bile canaliculi in different media conditions.

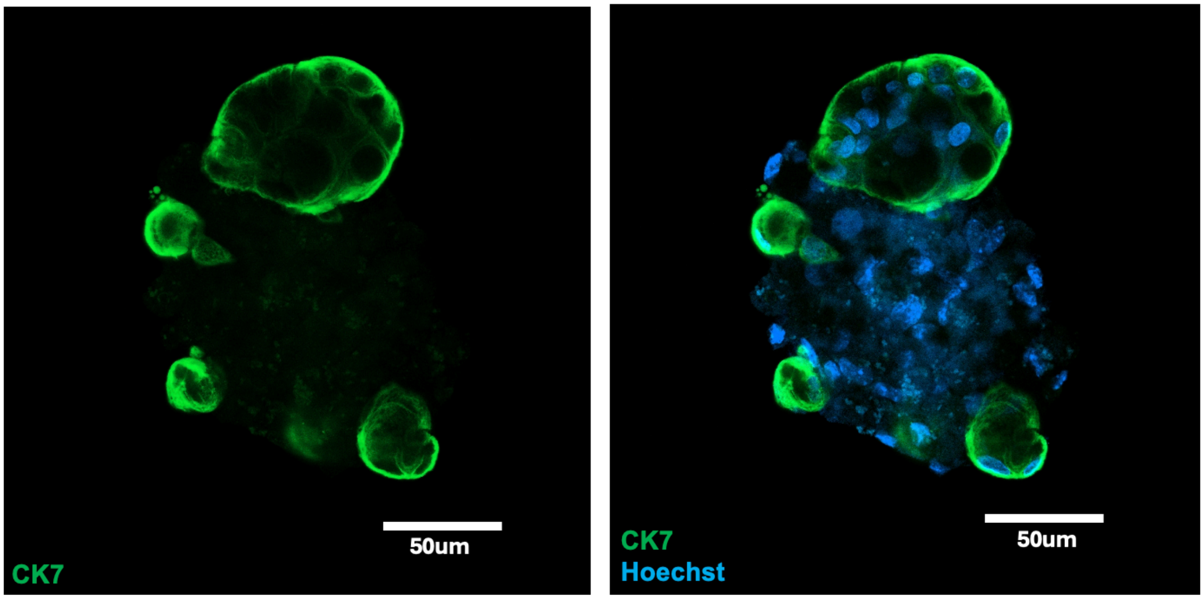


Supplementary Figure 5: CK7-positive ductule structures form within aHBOs.

Table S1: Media Formulations

| Hepatocyte Media | | |
|---|---|---|
| **Reagent** | **Vendor + Cat No.** | **Concentration** |
| DMEM with L-glutamine | Corning, 10-017-CV | N/A |
| ITS+ Premix Universal Culture Supplement | Corning, 354352 | 1% |
| Fetal bovine serum | GeminiBio | 10% |
| Dexamethasone | R&D Systems, 1126/100 | 0.04 ug/ml |
| Glucagon | Sigma-Aldrich, G2044-1MG | 7 ng/ml |
| HEPES | Sigma-Aldrich, H0887-20ML | 15mM |
| Penicillin/streptomycin | Life Technologies, 15140122 | 1% |

| ICO expansion media | | |
|---|---|---|
| **Reagent** | **Vendor + Cat No.** | **Concentration** |
| B27 Supplement 50x, minus vitamin A | Life Technologies, 12587-010 | 1x |
| N2 supplement 100x | Life Technologies, 17502-048 | 1x |
| N-acetylcysteine | Sigma-Aldrich, A0737-5MG | 1mM |
| Recombinant human [Leu15]-gastrin I | Sigma-Aldrich, G9145 | 10nM |
| Recombinant human EGF | Peprotech, AF-100-15 | 50ng/ml |
| Rspo1-conditioned medium | Made in house | 10% |
| Recombinant human FGF10 | Peprotech, 100-26 | 100ng/ml |
| Recombinant human HGF | Peprotech, 100-39 | 25ng/ml |
| Nicotinamide | Sigma-Aldrich, N0636 | 10mM |
| A83-01 | Tocris Bioscience, 2939 | 5uM |
| Forskolin | Enzo, BML-CN100-0010 | 10uM |
| Advanced DMEM/F12 | Thermo Fisher, 12634010 | N/A |

| Cholangiocyte media | | |
|---|---|---|
| **Reagent** | **Vendor + Cat No.** | **Concentration** |
| Nicotinamide | Sigma-Aldrich, N0636 | 10mM |
| Sodium bicarbonate | Sigma-Aldrich, S5761 | 17mM |
| 2-phospho-L-ascorbic acid trisodium salt | Sigma-Aldrich, 49752 | 200uM |
| Sodium pyruvate | Thermo Fisher, 11360070 | 0.63mM |
| Glucose | Invitrogen, 15023021 | 14mM |
| HEPES | Sigma-Aldrich, H0887-20ML | 20mM |
| ITS+ Premix Universal Culture Supplement | Corning, 354352 | 1% |
| Dexamethasone | R&D Systems, 1126/100 | 0.1uM |
| Glutamax | Lonza, BE17-605E/U1 | 1% |
| Penicillin/streptomycin | Life Technologies, 15140122 | 1% |
| Recombinant human EGF | Peprotech, AF-100-15 | 20ng/ml |
| Rspo1-conditioned medium | Made in house [15] | 10% |
| Recombinant human DKK1 | Abcam, ab155623 | 100ng/ml |
| William's E Medium, no phenol red | Invitrogen, A1217601 | N/A |

Table S2: List of qPCR primer sequences

| Gene Name | Forward Primer | Reverse Primer |
|---|---|---|
| HMBS | ACGGCTCAGATAGCATACAAGAG | GTTACGAGCAGTGATGCCTACC |
| BSEP | AGCCACACAGACCAGGATGTTG | CAATGAACCGCCTCTCCTTTCC |
| MRP2 | GCCAACTTGTGGCTGTGATAGG | ATCCAGGACTGCTGTGGGACAT |
| NTCP | GCTCTCTTCTGCCTCAATGGAC | AGTGGTCCAATGACTTCAGGTGG |
| CDH1 | GCCTCCTGAAAAGAGAGTGGAAG | TGGCAGTGTCTCTCCAAATCCG |
| CYP7A1 | CAAGCAAACACCATTCCAGCGAC | ATAGGATTGCCTTCCAAGCTGAC |
| KRT7 | TGTGGATGCTGCCTACATGAGC | AGCACCACAGATGTGTCGGAGA |
| KRT19 | AGCTAGAGGTGAAGATCCGCGA | GCAGGACAATCCTGGAGTTCTC |
| GGT1 | TGACGTACCACCGCATCGTAGA | CAGCGAAGAACTCGGAGGTCAT |
| CFTR | GGAGAGCATACCAGCAGTGACT | TTCCAAGGAGCCACAGCACAAC |
| EPCAM | GCCAGTGTACTTCAGTTGGTGC | CCCTTCAGGTTTTGCTCTTCTCC |
